# Estimating soil mineral nitrogen from data-sparse field experiments using crop model-guided machine learning approach

**DOI:** 10.1101/2024.09.03.610387

**Authors:** Rishabh Gupta, Satya K. Pothapragada, Weihuang Xu, Prateek Kumar Goel, Miguel A. Barrera, Mira S. Saldanha, Joel B. Harley, Kelly T. Morgan, Alina Zare, Lincoln Zotarelli

## Abstract

Sandy soils are susceptible to excessive nitrogen (N) leaching under intensive crop production which is linked with the soil’s low nutrient holding capacity and high-water infiltration rate. Estimating soil mineral nitrogen (SMN) at the daily time-step is crucial in providing fertilizer recommendations balancing plant nitrogen use efficiency (NUE) and N losses to the environment. Crop models [e.g., Decision Support System for Agrotechnology Transfer (DSSAT)] can simulate the trend of SMN in varied fertilizer rates and timing of application but are unable to replicate its magnitude due to the inability to capture high-water table conditions in a sub-irrigated soil. As an alternative to such physics-based model, time-series deep learning (DL) models based on a long short-term memory (LSTM) are promising in understanding nonlinearity among complex variables. Yet, purely data-driven DL models for crops are difficult to obtain due to the insufficient amount of data available and the excessive costs with producing more data. To address this challenge, a hybrid model (hybrid-LSTM) was developed by leveraging both the DSSAT and LSTM models to estimate daily SMN primarily using daily weather, applied fertilizer rates-timings, and the SMN sparse observations. This study used the observations from field trials conducted between 2010-2014 in Hastings, FL. The first step was to calibrate the DSSAT-SUBSTOR-Potato model to produce reliable SMN of the topsoil for treatments with varied N applied fertilizer rates split among the pre-planting, emergence, and tuber-initiation stages of the potato crop. Thereafter, the hybrid-LSTM model was trained on the calibrated DSSAT simulated SMN time-series and fine-tuned its predictions using the observed SMN to improve DSSAT simulated SMN. The hybrid-LSTM model was then tested on both calibrated and uncalibrated DSSAT SMN simulations where it outperformed the DSSAT model (range of improvement ranged ∼18-30% on comparing the normalized root mean squared error) in providing reliable estimates of SMN across most of the farms and years. This novel hybrid modeling approach could guide stakeholders and farmers to build sustainable N management with improved crop NUE and yield and help in minimizing environmental losses.

## 1. Introduction

Florida is a significant off-season contributor of fresh potatoes (*Solanum tuberosum L.*) consistently producing approximately 30% of the total spring production of the United States (NASS -Quick Stats, 2021). Most of the potatoes are grown in the northeastern region of the state, which is dominated by uncoated fine sand soil-type (USDA-NASS, 2019). These soils are highly vulnerable to excessive nitrogen (N) leaching due to rapid water infiltration (Zotarelli et al., 2007). Moreover, there is an impermeable soil layer below the surface, which allows subirrigation by maintaining a high-water table throughout the growing season (da Silva et al., 2018; Dukes et al., 2010). At the same time, it also poses challenges in effectively draining excess water during heavy rainfall. The field relies on irrigation furrows to distribute fresh ground water into the fields, connected to surrounding drainage ditches. Under heavy rainfall conditions, the water table level in the field is receded and excess water is moved off-site by gravity carrying soluble nutrients, especially N, from the top layer of the soil. Hence, multiple N fertilizer applications are required to compensate for the excess N leaching (Errebhi et al., 1998; Scholberg et al., 2013; Simonne et al., 2010).

Several studies have tried to estimate ideal fertilizer application rates and timings for potatoes in Florida (da Silva et al., 2023; Rens et al., 2018, 2016b, 2016a, 2015b, 2015a; Zotarelli et al., 2015, 2014). Typically, the N fertilizer applications should synchronize with the potato growth and N uptake. The N uptake rates are higher between mid-vegetative growth to mid-tuber initiation; hence, N applications between emergence and tuber-initiation are crucial to maximize tuber yield (Rens et al., 2018, 2016a; Zotarelli et al., 2015). Zotarelli et al. (2015) and Rens et al. (2018) reported that N application at pre-planting could be highly prone to leaching particularly in cases of heavy rainfall projections before plant emergence. Hence, they recommended applying N fertilizer close to planting rather than at pre-planting. Another finding by Zotarelli et al. (2014) suggests keeping N application rates 224-280 kg-N ha^-1^ to maximize tuber yield under heavy rainfall conditions. Rens et al. (2018) found that tuber yield peaked when N application specifically at emergence varied from 128-168 kg-N ha^-1^ under the same circumstances. Similarly, N application at tuber-initiation was observed to be crucial with ∼62% N use efficiency and found to have no significant increase in tuber yield when applied above 56 kg-N ha^-1^ (Rens et al., 2016a; Zotarelli et al., 2014). Therefore, it becomes crucial to adaptively manage fertilizer application rates and timings in highly uncertain rainfall conditions to minimize N leaching and maximize the potato yield and N use efficiency.

Estimating SMN at a higher frequency is crucial for making informed decisions for fertilizer applications to aid growers and stakeholders in improving tuber yield/N use efficiency without harming the environment. However, in-situ measurement of SMN is time-consuming, labor-intensive, and highly expensive. Consequently, there is a need to develop a reliable framework and methodology to estimate SMN at a higher frequency using the limited experimental data. Previous research employed process-based cropping system models, such as DSSAT (Hoogenboom et al., 2019; Jones et al., 2003), Agricultural Production Systems sIMulator (APSIM) (Holzworth et al., 2014; Keating et al., 2003), and other agroecological models to replicate the interaction between soil, water, and crop under different environmental and geographical setting (Gupta et al., 2022; Liu et al., 2011; Moriasi et al., 2013; Salmerón et al., 2014; Saseendran et al., 2007). However, these models may not capture the variability of SMN in the sandy soils. This variability is particularly difficult to capture when caused by uncertain rainfall events and water table fluctuations as these models could not simulate three-dimensional movement of N dynamics in the soil layers, necessary for precise estimation of SMN (Archontoulis et al., 2014; Raymundo et al., 2017; Wallach and Thorburn, 2014). Hence, these models may not provide accurate estimates of SMN required for narrowing weather-specific crop nutrient recommendations. Additionally, calibrating these models could be tedious and time-consuming as it would require fine-tuning the extensive list of parameters (Akhavizadegan et al., 2021; Seidel et al., 2018). Furthermore, one would need to re-calibrate these models for different climate and soil conditions. Therefore, current circumstances call for devising new approaches and modeling frameworks that possess transferable learning, reducing the need for repeated calibration.

Deep learning (DL) techniques have been extensively embraced by researchers for interpreting intricate agroecological systems, as they could effectively learn complex relationships between plants, soil, and climate (Kamilaris and Prenafeta-Boldú, 2018). A significant advantage of DL is their transfer learning ability to adapt to completely new scenarios (Weiss et al., 2016), such as, for different crop, soil, or climate conditions. Most research in the field of agriculture used remote-sensing data (multispectral/hyperspectral dataset or satellite imageries) for building DL models to estimate the crop growth stages (Wang et al., 2022; Yue et al., 2020), detect pests/diseases (Hadipour-Rokni et al., 2023; Mohanty et al., 2016), predict leaf area indices (Ilniyaz et al., 2023), soil properties (Padarian et al., 2019; Zhang et al., 2022), and evapotranspiration (Sharma et al., 2022).A deep recurrent neural network-based Long Short-Term Memory (LSTM) (Hochreiter and Schmidhuber, 1997) has significant popularity and proven its effectiveness in capturing the temporal variations in the high-frequency time series data in the agricultural domain (Datta and Faroughi, 2023; Gauch et al., 2021; Zhang et al., 2022). For instance, Saha et al. (2023) built an LSTM model to estimate daily stream nitrate concentration for 42 sites in data-sparse watersheds in Iowa, U.S.A., with a Nash-Sutcliffe efficiency of ∼75%. Similarly, Datta and Faroughi (2023) successfully predicted the soil moisture content for the next hour, a day, a week, and a month in advance, achieving a R-squared value of ∼0.95, developing a multiheaded-LSTM model, training the model using 15-minute interval field measurements.

Nevertheless, time-series machine learning (ML) models like LSTM necessitate daily, or at least, discrete long-term observations of SMN at a higher frequency, to effectively learn the temporal variation caused by weather, irrigation, and N applications (Hua et al., 2019). Building an effective time-series ML model like LSTM with limited observations (5-6 sampling per season) could not be possible. Hence, the DSSAT model was employed to accompany the LSTM model to accurately predict the response of weather and N application (rates/timings) on daily SMN values during the crop cycle (Fig. 1). Numerous studies have affirmed that ML could serve as a companion to crop models rather than a competitor (Everingham et al., 2016; van Klompenburg et al., 2020; Zhang et al., 2023). Feng et al. (2020) asserted that comprising ML-based models with biophysical agroecosystem models has the potential to provide an improved evaluation of the changing climate on wheat yield in Australia. Similarly, Shahhosseini et al. (2021) found promising results in adding APSIM model simulated features in the ML models for corn yield prediction in the U.S. Corn Belt. In the present study, we explored a similar approach to develop a hybrid model [DSSAT + LSTM = hybrid-LSTM] to estimate the SMN between the soil sampling dates for improved recommendations on crop nutrient management. Initially, we calibrated the DSSAT model to simulate daily SMN. Afterward, the DSSAT simulated daily SMN values were used to train the hybrid-LSTM model. Later, we fine-tuned the hyperparameters of hybrid-LSTM using limited sampled in-situ SMN observations to improve the SMN estimates. The overall goal of this integration was to replicate the SMN reducing the need for tedious calibration of the DSSAT model by leveraging its strengths (ability to simulate dynamics of the cropping system) while mitigating its weaknesses (inability to capture soil nutrients and water dynamics in a sub-irrigated high water-table conditioned cropping systems) via LSTM modeling. The objective of this study was to develop the hybrid-LSTM model that accurately estimate daily SMN throughout a potato crop growing season using limited SMN data. The outcomes of this study can be further used to support informed decisions on fertilizer application rates, timings to improve the tuber yield and reduce the environmental impacts.

**Figure 1.**
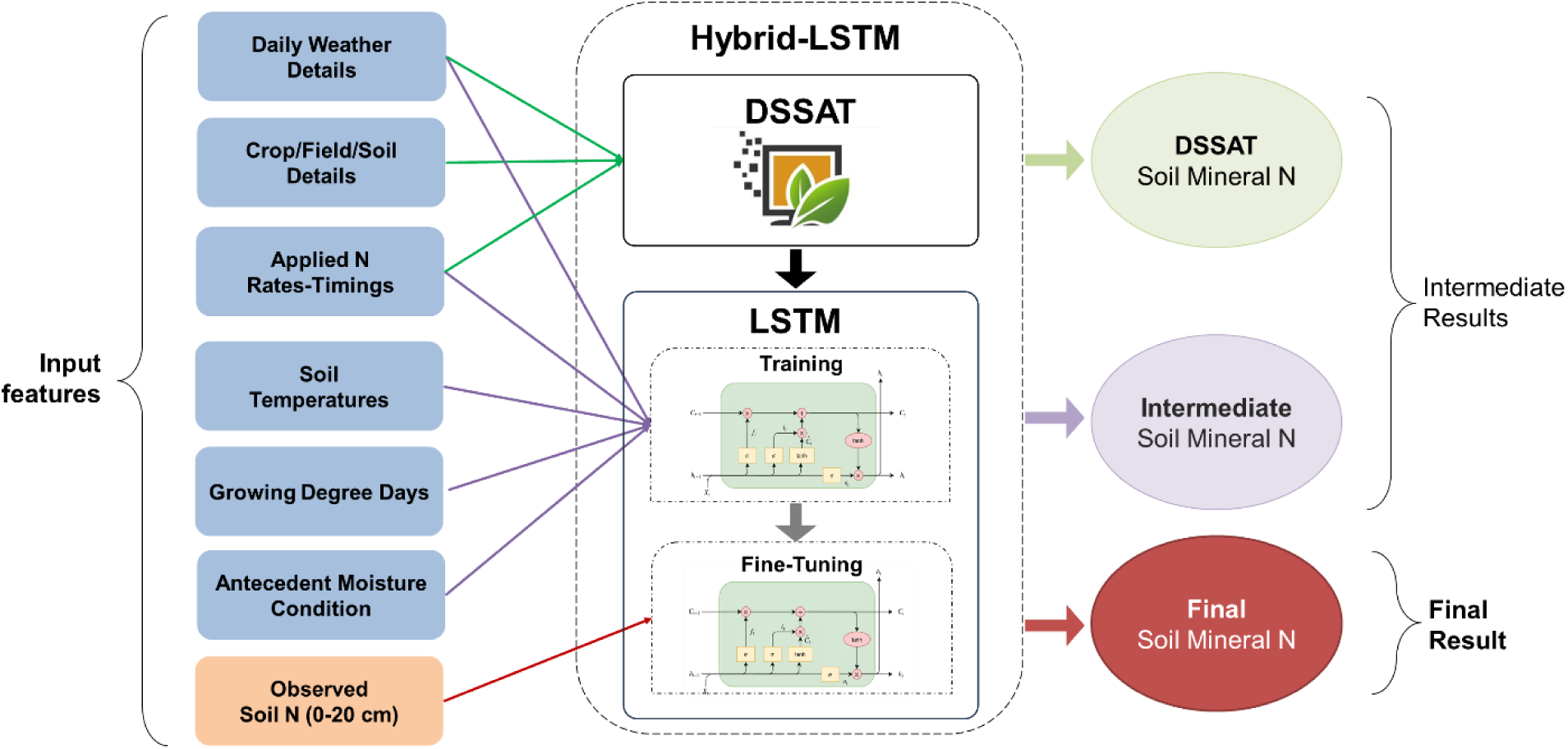
Block diagram to demonstrate the integration of two models (DSSAT and LSTM), input features, and the workflow of the hybrid-LSTM model.

## 2. Materials and Methods

### 2.1. Field experiments overview

A series of field experiments were conducted on several commercial potato fields in Hastings, FL, U.S. from 2011-2014. Experiments carried out in the spring of 2011 and 2012 used the ‘FL 1867’ potato cultivar in Farm 1 (F1), Farm 2 (F2), and Farm 3 (F3), and the ‘Atlantic’ potato cultivar in Farm 4 (F4). Field trials conducted in 2013 and 2014, used the ‘FL 1867’ potato cultivar in F1 and the ‘Atlantic’ potato cultivar in F4. Each experiment was comprised of eight treatments combining three application timings and N fertilizer rates (Table 1). All the treatments of 2011-2012 experiments received a fixed application (56 kg-N ha^-1^) of granular ammonium nitrate during pre-plant followed by the combination of four different rates of liquid urea ammonium nitrate (0, 56, 112, and 168 kg-N ha^-1^) at the plant emergence and two different rates of liquid urea ammonium nitrate (56, 112 kg-N ha^-1^) as side-dress at the tuber initiation. The treatments of 2013-2014 experiments received N fertilizer rates of 0 and 56 kg-N ha^-1^ at the pre-plant followed by N rates of 0, 56, 112, and 168 kg-N ha^-1^ at the plant emergence and a fixed rate of 56 kg-N ha^-1^ at the tuber initiation. All the treatments in the last two years of experiments received granular ammonium nitrate. The predominant soil types in F1, F2, F3, and F4 are Ultic Alaquod, Arenic Glossaqualf, Arenic Endoaqualf, and Aeric Alaquod, respectively. All the field trails from 2011-2014 were conducted under seepage irrigated maintaining the water table at around 55 cm below the top of the raised soil bed for potato planting. More details of the experimental design and results can be found in (Rens et al., 2018, 2015a; Zotarelli et al., 2015, 2014).

**Table 1.** Nitrogen fertilizer application rates and timings (at pre-planting, plant emergence, and tuber initiation) along with total N applied during potato season for the two set of field experiments. One set of experiment was conducted in 2011 and 2012, while other set of experiment was conducted in 2013 and 2014.

| Experiment Year | Treatments | N fertilizer rate applied (kg-N ha<sup>-1</sup>) at |  |  | Total N applied |
| --- | --- | --- | --- | --- | --- |
|  |  | Pre-planting | Plant Emergence | Tuber Initiation |  |
| 2011 and 2012<br>(F1, F2, F3, F4) | 1 | 56 | 0 | 56 | 112 |
|  | 2 | 56 | 0 | 112 | 168 |
|  | 3 | 56 | 56 | 56 | 168 |
|  | 4 | 56 | 56 | 112 | 224 |
|  | 5 | 56 | 112 | 56 | 224 |
|  | 6 | 56 | 112 | 112 | 280 |
|  | 7 | 56 | 168 | 56 | 280 |
|  | 8 | 56 | 168 | 112 | 280 |
| 2013 and 2014<br>(F1 and F4) | 1 | 0 | 0 | 56 | 56 |
|  | 2 | 56 | 0 | 56 | 112 |
|  | 3 | 0 | 56 | 56 | 112 |
|  | 4 | 56 | 56 | 56 | 168 |
|  | 5 | 0 | 112 | 56 | 168 |
|  | 6 | 56 | 112 | 56 | 224 |
|  | 7 | 0 | 168 | 56 | 224 |
|  | 8 | 56 | 168 | 56 | 280 |

### 2.2. Data description and preparation

This study uses weather-soil-crop management information to use the DSSAT and ML models for estimating daily SMN. Daily weather data of Hastings, FL (29.6933^°^N, 81.4449^°^W), comprised of rainfall, average, minimum, maximum air temperatures (at 60 cm height), solar radiation, dew point, relative humidity, and wind speed, was collected from the Florida Automated Weather Network data access portal (FAWN, 2022) from 2010-2014. Daily soil temperatures (average, minimum, maximum) were also collected from the FAWN data access portal for the same period. Field management information such as potato planting/emergence/tuber initiation/harvesting dates, details of fertilizer application (rates and timings), initial soil condition, previous cropping history, and other relevant information was collected from the earlier studies (mentioned in section 2.1). Soil surface information, such as soil albedo, slope, drainage class, runoff potential, and soil layered information, for instance, soil wilting point and field capacity, saturated water content, bulk density, organic matter content, and soil root growth factor were collected from previous studies and Soil Survey Geographic Database (da Silva et al., 2018; Rens et al., 2022; Reyes-Cabrera et al., 2016; SSURGO Database, 2018). The DSSAT model was set up using the weather-soil-crop information collected from various sources by creating its weather, soil, and experiment files. The observed datasets from the field trials such as SMN, potato aboveground biomass/N accumulation, and tuber weight (dry and fresh)/N accumulation was used to evaluate the performance of the DSSAT model for replicating the potato cropping system.

Likewise, daily weather information (rainfall, average air temperature), soil average temperature, applied N fertilizer rate-timing, and the DSSAT simulated daily SMN were considered in developing the hybrid-LSTM model. Later, all the dataset was arranged into different samples, where every sample corresponds to a specific farm and a specific year with all the N applied rates-timings treatments. Every sample contained training input of a specific farm-year (farm-year represents the data of a specific farm and a specific year), which includes data for all the input features and a target vector for the same. Since the hybrid-LSTM was first trained with the DSSAT simulated data and then with the observed SMN, we had two sets of training data, the difference being the number of data points per sample. The first training set with DSSAT simulated data had daily data of all the features for the whole crop cycle, while the second training set contained only those points when the SMN was sampled at the experiment site for fine-tuning the model (more detail provided in section 2.4.2).

### 2.3. Model description

#### 2.3.1. DSSAT model

The process-based cropping system model DSSAT v4.8 can simulate soil water-nutrient dynamics and the growth and development of more than 42 crops (Hoogenboom et al., 2023, 2019; Jones et al., 2003). It employs the SUBSTOR-Potato model (Griffin et al., 1993; Ritchie et al., 1995) of DSSAT to simulate daily phenological development, biomass accumulation, and tuber yield in a climate-soil varied setting. SUBSTOR-Potato model of DSSAT has five cultivar-type parameters-G2 controls leaf area expansion after tuber initiation (cm^2^ m^-2^ d^-1^), G3 controls potential tuber growth rate (g m^-2^ d^-1^), PD suppresses tuber growth after tuber induction (relative index, dimensionless), P2 controls tuber initiation sensitivity to long photoperiod (relative index, dimensionless), and TC is upper critical temperature for tuber initiation (^°^C). It also has two radiation use efficiency (RUE) parameters-RUE1 (3.5 g plant dry matter per MJ photosynthetically active radiation) and RUE2 (4.0 g plant dry matter per MJ photosynthetically active radiation) to represent before and after tuber initiation. We used Ritchie water balance, FAO-56, soil conservation service, and Suleiman-Ritchie methods to estimate soil hydrology, evapotranspiration, soil infiltration, and soil evaporation, respectively (Ritchie et al., 1998). Active, intermediate, and passive soil organic matter pools were determined using the CENTURY model in DSSAT (Gijsman et al., 2002).

The DSSAT model was calibrated for plant/tuber biomass/N accumulation and soil mineral nitrogen. The soil parameters-wilting point, field capacity, saturated water content, bulk density, organic matter content, soil root growth factor, and initial soil conditions were adjusted carefully to closely match with the real field conditions based on previous literatures (da Silva et al., 2018; Rens et al., 2022; Reyes-Cabrera et al., 2016) and Soil Survey Geographical Database (SSURGO Database, 2018) to simulate SMN for the topsoil layer (Table 2). Genotype coefficients of the potato cultivar were calibrated to replicate tuber/plant biomass and N accumulation (Table 3). Other parameters, for example, sprout length and irrigation depth were also found sensitive to plant/tuber growth, hence these parameters were also adjusted accordingly for modeling purposes. The DSSAT model was calibrated using field observations of four farms (F1-F4) from 2011-2012; while the model was evaluated using field observations of two farms (F1, F2) from 2013-2014.

**Table 2.** Soil layered parameters were used for all the farms to calibrate the DSSAT model. Source: (da Silva et al., 2018; Rens et al., 2022; Reyes-Cabrera et al., 2016; SSURGO Database, 2018).

| Soil Depth (cm) | Wilting Point (cm<sup>3</sup>cm<sup>-3</sup>) | Field Capacity (cm<sup>3</sup>cm<sup>-3</sup>) | Saturated water content (cm<sup>3</sup>cm<sup>-3</sup>) | Root growth factor (cm<sup>3</sup>cm<sup>-3</sup>) | Bulk Density (g cm<sup>-3</sup>) | Organic Carbon (%) |
| --- | --- | --- | --- | --- | --- | --- |
| 5 | 0.055 | 0.128 | 0.456 | 1.00 | 1.53 | 0.65 |
| 15 | 0.055 | 0.125 | 0.456 | 1.00 | 1.53 | 0.65 |
| 30 | 0.047 | 0.123 | 0.429 | 0.44 | 1.54 | 0.63 |
| 45 | 0.036 | 0.122 | 0.419 | 0.02 | 1.57 | 0.44 |
| 60 | 0.036 | 0.119 | 0.419 | 0.01 | 1.57 | 0.44 |
| 90 | 0.036 | 0.110 | 0.407 | 0.00 | 1.61 | 0.18 |
| 120 | 0.036 | 0.110 | 0.407 | 0.00 | 1.61 | 0.18 |
| 150 | 0.033 | 0.108 | 0.401 | 0.00 | 1.63 | 0.08 |

**Table 3.** Calibrated genotype coefficients to simulate the growth and development of ‘Atlantic’ and ‘FL1867’ potato cultivar.

| Coefficients | Definitions | Values |
| --- | --- | --- |
| G2 | Leaf area expansion rate after tuber initiation (cm<sup>2</sup> m<sup>-2</sup> d) | 1000 |
| G3 | Potential tuber growth rate (g m<sup>-2</sup> d) | 25.0 |
| PD | Index that suppresses tuber growth during the period that immediately follows tuber induction | 0.9 |
| P2 | Tuber initiation sensitivity to long photoperiods | 0.6 |
| TC | Upper critical temperature for tuber initiation (°C) | 17.0 |

#### 2.3.2. LSTM model

A long short-term memory (LSTM) is a type of recurrent neural network designed to process sequential (or time-series) data by effectively capturing long-term dependencies (Hochreiter and Schmidhuber, 1997). The structure of each LSTM cell is the fundamental block of the network. It consists of a memory state along with a gating mechanism to selectively store, forget, and process the output information (Staudemeyer and Morris, 2019). Whenever a new input sequence is passed through an LSTM cell, the input and forget gates within its cell determine what information is needed to be stored in and forgotten from the cell state, respectively, at a particular timestep. The output gate processes information from the cell state, which is the output of LSTM at the consecutive time step. This process is repeated for each time step to produce a time series output. Sequence length is one of the crucial parameters of LSTM architecture. The sequence length refers to the number of time steps or observations considered as input to predict the consecutive time step (Hua et al., 2019). It affects the computational complexity, memory requirements, and the model’s ability to capture long-term dependencies. While longer sequences may lead to vanishing gradients, overfitting, and increased computational demands, shorter sequences offer computational efficiency at the cost of reduced context (Hua et al., 2019). Every input that goes into the LSTM network is a sequence of data which is represented as an input feature matrix and a target vector, fed into the model one after the other while training.

#### 2.3.3. The hybrid-LSTM model – An integration of the DSSAT and LSTM model

The main goal of the hybrid-LSTM model development was to estimate the daily time-series of the SMN using weather, soil and air temperatures, and fertilizer applications (rates, timings). However, training a ML model exclusively on experimental SMN data was infeasible due to the lack of observed data available (only 5-6 soil samplings per crop cycle per experiment) to effectively capture the complex relationships and estimate daily SMN. Hence, the DSSAT model was used, which could simulate daily SMN throughout the crop cycle along with other input features to train a time-series (LSTM) model. However, the current version of DSSAT (v4.8) is unable to replicate a sub-irrigated cropping system in a high-water table condition. Hence, the study was aimed to improve over DSSAT simulated SMN in such data-sparse, sub-irrigated, and high-water table conditioned potato cropping system in northeastern Florida using hybrid-LSTM model. The key concept was to train a time-series ML model (LSTM) on the DSSAT simulated SMN to learn the interaction between the features and the simulated SMN. The ML model (LSTM) then trained again with these features but with experimental SMN on those days experimental data is available, which is called as fine-tuning process. This integration of DSSAT and LSTM was called the hybrid-LSTM model for ease of explanation.

### 2.4. Setup and training of the hybrid-LSTM model

#### 2.4.1. Feature Engineering and Parameterization

Identifying the right features for the ML model, known as feature engineering, is crucial for efficient model training. As a part of the study, a comprehensive list of potential features was prepared to subsequently conduct a feature importance test, identifying the features that exhibited a strong correlation with the experimental SMN values. The comprehensive list of potential features included daily rainfall, minimum-maximum-average air and soil temperatures, evapotranspiration, and applied N fertilizer rate-timing. The list also included the DSSAT simulated data-daily SMN and plant N, in the feature selection process. Furthermore, a feature to capture the interaction of rainfall and applied N fertilizer rates (multiplying their normalized values only when rainfall occurred on the N application day) was also included in the potential features list to supplement the model understanding of potential N leaching. We also considered features such as antecedent soil moisture condition (it was assumed dry when last 5 days of rainfall accumulation was less than 35 mm, and wet in vice versa) (Gray et al., 1982) and growing degree days of potato crop (Hristine et al., 2005). After performing feature importance testing, the selected features were rainfall, average air temperatures, soil average temperatures, applied N fertilizer rate-timing, rainfall and applied N fertilizer rate interaction feature, and the DSSAT simulated SMN.

The sequence length of the hybrid-LSTM model was optimized to 10 days by conducting a grid search through experimental runs to effectively capture meaningful patterns between input features and the target variable. The hidden units and the number of layers were chosen such that the total number of the hybrid-LSTM parameters did not exceed the total number of input data points, minimizing the model complexity. A specific optimizer and learning rates were used for training and fine-tuning the hybrid-LSTM model. The L-2 regularization was applied to the weights of the hybrid-LSTM model to smoothen the output and avoiding unwarranted peaks and valleys in the estimated SMN curve. Moreover, air and soil temperatures were detrended (to remove seasonality from winter to summer) to avoid any temporal dependencies in the hybrid-LSTM model.

#### 2.4.2. Training Procedure

The training procedure was divided into two parts. The first part included the training of the hybrid-LSTM model on the calibrated DSSAT simulations of SMN. This part of the training was to understand the relationship between input features and DSSAT simulated SMN. The second part of the training procedure involved refining the SMN estimated by the hybrid-LSTM model further using sparsely available observed SMN. The developed hybrid-LSTM model’s performance was tested against SMN observations on the sampling days while feeding the unseen-calibrated and - uncalibrated DSSAT simulations of different farm-year combinations in the hybrid-LSTM model.

Since the soil parameters of the DSSAT model were not adjusted for specific farm during calibration, assuming similar soil properties among all the farms, the performance and trend of the DSSAT simulated SMN was not consistent for all the farms-years compared to the observed SMN. Hence, the farms-years were qualitatively classified into “good”, “bad”, and “uncertain” categories based on the agreement between the DSSAT-simulated and observed SMN trends before the model training. This classification was performed based on the visual interpretation as it was difficult to conform to a specific evaluation metric that can compare trends between two quantities which are not distributed on the similarly over time (e.g., DSSAT simulated SMN is continuous, daily data is available, but observed SMN is sparsely distributed over time). The good farms exhibited consistent trends, while the bad farms showed contrasting patterns. The uncertain farms had incomplete agreement across all sampling stages. Thereafter, different sets of training and testing with different farms-years combinations were generated at random, emphasizing more on good farms-years followed by bad and uncertain farms-years. This process allowed us to assess the model’s ability to handle variations introduced by challenging farms-years (bad and uncertain farms-years) simulations for effective learning. By incorporating this approach, we ensured that the hybrid-LSTM model effectively captured the complexities of different categories (good/bad/uncertain), leading to accurate predictions of SMN. For every combination, the hybrid-LSTM model was trained several times, incorporating a cross-validation approach for each repetition. Finally, the best performing set of training and testing farms-years was selected using the performance metrics mentioned in section 2.5. The need of multiple iterations for training the hybrid-LSTM model stemmed due to the SMN observations scarcity, posing a formidable challenge in estimating a precise SMN line curve for each crop cycle. Conducting multiple training iterations not only reduced the noise from the line curve but also enabled the representation of daily SMN estimated with an area curve. The SMN area curve was derived using the standard deviation across all the iterations from the most successful set of training and testing farms-years, while the average of all the iterations was used to create the smooth line curve for SMN.

The final set of farms-years used for training are F1-2011, F2-2011, F3-2011, F2-2012, F3-2012, F4-2012, F4-2013, and F4-2014. The testing farms are F4-2011, F1-2012, F1-2013, and F1-2014 for all treatments (where F2-2012 and F4-2013 were uncertain farms-years, and F1-2012 and F4-2012 were bad farms-years, rest are good farms-years).

### 2.5. Statistical approaches for the model evaluation

The hybrid-LSTM and DSSAT model performances were assessed by comparing the estimated or simulated SMN line and area curves from these models against the scarce observations. Since there were four replicates of the SMN observed and we estimated the SMN line/area curve using the hybrid-LSTM and DSSAT models, it was challenging to evaluate the model performance by comparing the line/area curve with the observed replicates. Hence a metric was formulated to evaluate the model estimated SMN line/area curve considering the observed replicant bounds of SMN. The metric calculates the normalized absolute error using the lower and upper bounds of replicates given if the SMN predicted values at the sampling stages were closer to the lower and upper bounds of replicates, respectively (Eq. 1). This metric was called the passing error (*PE*). *PE* = 0 would be an ideal case when the estimated SMN line curve passed through the replicant bounds of the observed SMN on all the sampling dates. The frequency of such alignment across all sampling dates was noted in all the farms-years. This metric was used to compare the performance of the hybrid-LSTM and the DSSAT models.

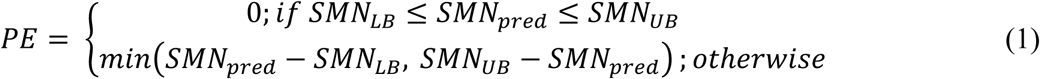

Other statistical performance metrics-normalized-root mean square error (*nRMSE*), and coefficient of determination (*R^2^*) were used to assess the simulated results from the DSSAT and the hybrid-LSTM models (Laperre et al., 2020; Moriasi et al., 2007). Due to the inadequate observed data (such as tuber and plant dry weights and N accumulation, tuber fresh weight, and SMN), the simulated outputs were also evaluated based on visual interpretations (randomly picking graphs and manually checking if the simulated/estimated values aligned with observed data) to ensure the coherence of the model.

## 3. Results and Discussion

The hybrid-LSTM model, an integration of a process-based crop simulation model-DSSAT and a time-series ML model-LSTM, was able to learn the complexity between the input features (e.g., rainfall, average air temperature, average soil temperature, and applied N rates) and the temporal variability in the SMN. The generalized hybrid-LSTM model trained on various experimental sites in Hastings, FL was able to predict the SMN daily values with the improvement of 22.2% (*nRMSE*) and 27.2% (*PE*) compared to the DSSAT simulated SMN estimates across all the farms and years. These improvements indicate the effectiveness of the hybrid modeling approach used in this study. In the subsequent sections, the detailed performance analysis of the DSSAT model (calibration and evaluation) and the hybrid-LSTM model (training and testing) is provided.

### 3.1. DSSAT model performance

#### 3.1.1. Potato growth simulation by DSSAT

To evaluate the performance of the SUBSTOR-potato model of DSSAT for potato growth and development, the simulated results on tuber dry weight (DW), plant DW, tuber N, plant N, and tuber yield fresh weight (FW) were compared against observed values (Fig. 2). The *R^2^* and *nRMSE* values for plant/tuber growth/N ranged from 0.75-0.92 and 26.3-32.0% during the DSSAT model calibration. The DSSAT model performance during the evaluation was also similar to calibration with the *R^2^* and *nRMSE* values for plant/tuber growth/N ranging from 0.51-0.92 and 29.1-51.3%, respectively. However, the DSSAT model underperformed in simulating tuber yield during calibration (*R^2^*=-0.08, *nRMSE*=14.6%), caused specifically due to consistent poor performance in all four farms in 2011. The DSSAT model underestimated the tuber yield in 2011 (average observed yield=46.2 Mg ha^-1^, average simulated yield=23.9 Mg ha^-1^). In contrast, the DSSAT model performance was satisfactory (*R^2^*=0.56, *nRMSE*=8.1%) during evaluation in 2013 and 2014 for two farms (F1, F2). Raymundo et al. (2017) also simulated similar results using the SUBSTOR-potato model of DSSAT with *nRMSE* of 21.0%, 37.2%, 85.3%, 40.4%, and 86.3%, respectively, for tuber FW, tuber DW, aboveground DW, tuber N update, and above ground N. Similarly, Wang et al. (2023) observed *nRMSE* ranging between 12.3 and 69.7% when simulating tuber DW using the same model from a two-year experiment in China with different irrigation and fertigation levels.

**Figure 2.**
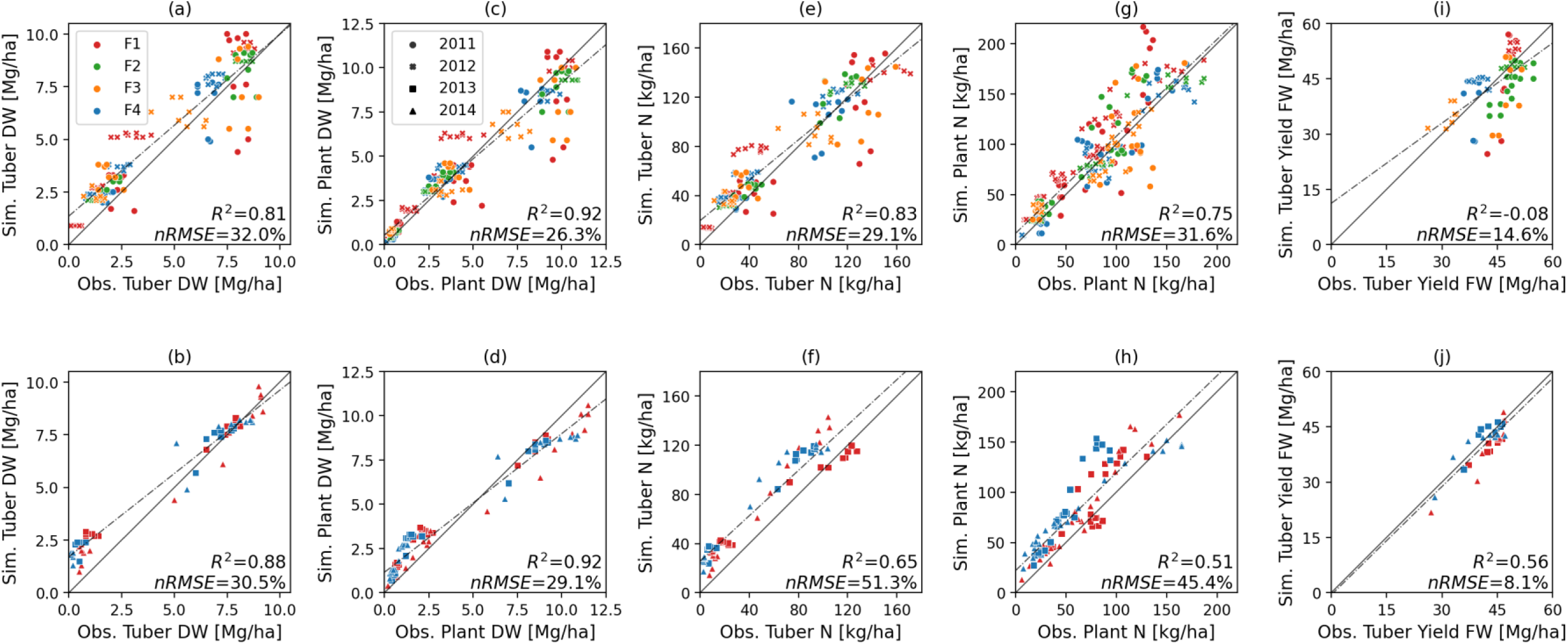
Regression plots comparing the observed and DSSAT – SUBSTOR Potato model simulated (a, b) tuber dry weight (DW), (c, d) plant DW, (e, f) tuber N, (g, h) plant N, and (i, j) tuber yield fresh weight (FW) while calibrating (first row, model calibrated using observations from four farms for 2011 and 2012) and evaluating (second row, model evaluated using observations from two farms for 2013 and 2014) the DSSAT model. [Note: Different colors of markers represent different farms whereas different shapes of markers represent different years, *R^2^* = Coefficient of Determination, *nRMSE* = Normalized root mean squared error].

A possible reason behind the DSSAT model’s poor performance in 2011 could be attributed to the atypical rainfall distribution in that year (Fig. 3). The 2011 season received less rainfall (80 mm, average of all farms) compared to other years (130, 401, and 145 mm in 2012, 2013, and 2014, respectively) after tuber-initiation when there is a rapid increase in the plant N uptake due to tuber bulking. The reduced rainfall after tuber initiation in 2011 and the upward soil water flux from the subirrigation could have induced less N leaching from the topsoil compared to other years. Moreover, poor drainage conditions due to an impermeable layer below the surface require minimal irrigation to maintain a high-water table in dry conditions. In contrast, the current version of the DSSAT model could not replicate such a complex high-water table system in the sub-irrigated potato fields. Moreover, the model treats the subsurface irrigation as surface irrigation, and hence, the model might have overestimated the N leaching by applying more than needed irrigation water amid dry conditions (not considering the impermeable layer to hold the water table level) which could have underestimated the simulated tuber yield. Overall, the DSSAT performance in simulating the potato growth was satisfactory well.

**Figure 3.**
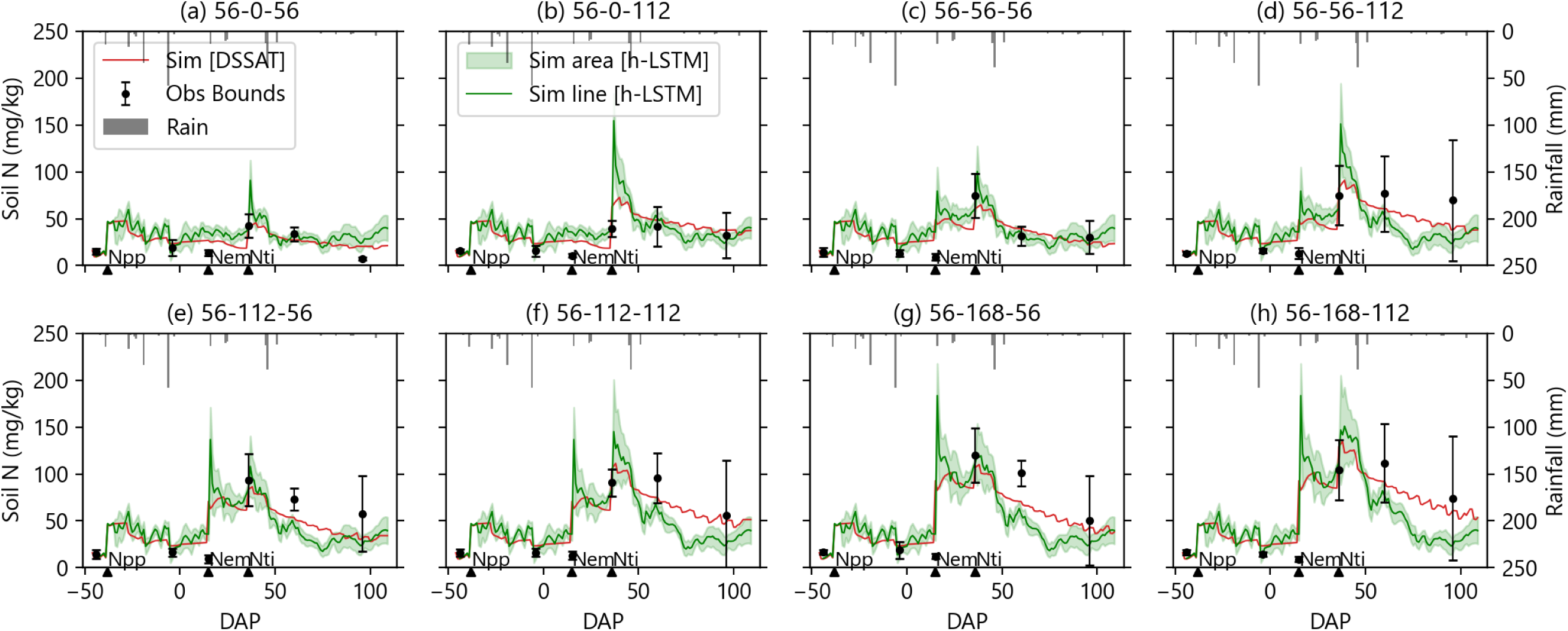
Comparison of the observed (black error bars, error bar represents the standard deviation within the replicates), DSSAT simulated (red line) and hybrid-LSTM estimated (green line, green area curve) soil N concentration (0-15 cm) in F3-2011 for different fertilizer N rates and timing of application treatments (subplot title of (a) is 56-0-56 which means N_pp_ =56 kg-N ha^-1^, N_em_=0 kg-N ha^-1^, and N_ti_ = 56 kg-N ha^-1^, the solid triangles below x-axis are the days after planting (DAP) when N fertilizer was applied for N_pp_, N_em_, and N_ti_) while training the hybrid-LSTM model [Sim = Simulated, Obs = Observed; N_pp,_ N_em_, and N_ti_ = N fertilizer applied at planting, emergence, and tuber initiation, respectively]

#### 3.1.2. Soil mineral nitrogen simulation by DSSAT

Figure 3 and 4 show the comparison of the DSSAT simulated and observed SMN values to illustrate the model performance during calibration and evaluation on F3-2011 and F1-2013, respectively, for all the N fertilizer rate and timing treatments. The DSSAT model performance was satisfactory during calibration and evaluation for simulating the daily SMN values. The *nRMSE* and *PE* for the observed vs. DSSAT simulated SMN values were 16.0% and 8.0%, respectively in calibration (Table 4). Whereas the *nRMSE* and *PE* for the observed vs. DSSAT simulated SMN values were 12.8% and 9.3%, respectively in the model evaluation (Table 4). These results reveled an improvement in simulating SMN (nitrate-N and ammonium-N) over previous study by Raymundo et al. (2017) who observed *nRMSE* of 95.5% and 140.1% using the earlier version of the DSSAT model. Overall, the current version of DSSAT model (calibrated in this study) was able to simulate the daily variation of SMN, however, not the order of magnitude of variation of SMN. For example, in Fig. 3, the DSSAT simulated SMN curve followed the pattern closely in most of the treatments (Fig. 3). However, DSSAT simulated SMN values were consistently underestimated in all the treatments. The possible reason for such performance of the DSSAT model could be attributed to its inability to replicate SMN in a high-water table condition maintained by the sub-irrigation of the fields. The daily high-water table level oscillation might have influenced the dynamics of water and N movement in the sandy soil in the observed conditions (da Silva et al., 2018). In contrast, the DSSAT model performance in simulating the SMN trend was reasonably well in heavy and more uniformly distributed rainfall conditions (Fig. 3, 4). For instance, the rainfall was more frequent and heavier after the tuber initiation (compared to before the tuber initiation) in F1-2013, and correspondingly, the DSSAT simulated SMN values also followed the same trend with observed SMN values after the tuber initiation N application (Fig. 4). This could be attributed due to limited need of irrigation in such conditions and the irrigation furrow would have been kept open to drain the excessive rainfall water. Overall, the DSSAT model was able to simulate the daily SMN values satisfactorily well given the continuous oscillation of the water table level and the presence of the sub-surface irrigation which is not supported as of the current version of the DSSAT model.

**Figure 4.**
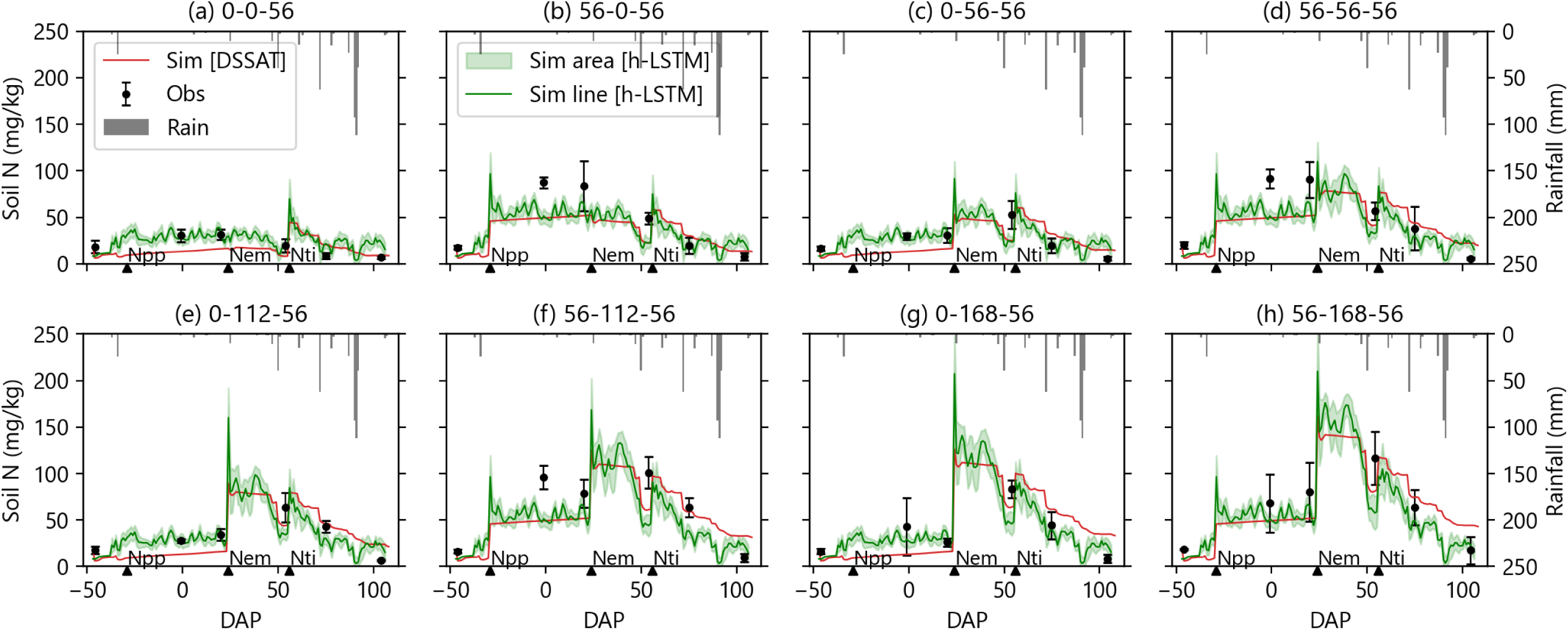
Comparison of the observed (black error bars, error bar represents the standard deviation within the replicates), DSSAT simulated (red line) and hybrid-LSTM estimated (green line, green area curve) soil N concentration (0-15 cm) in F1-2013 for different fertilizer N rates and timing of application treatments (subplot title of (a) is 0-0-56 which means N_pp_=0 kg-N ha^-1^, N_em_=0 kg-N ha^-1^, and N_ti_=56 kg-N ha^-1^, the solid triangles below x-axis are the days after planting (DAP) when N fertilizer was applied for N_pp_, N_em_, and N_ti_) while testing the hybrid-LSTM model [Sim = Simulated, Obs = Observed; N_pp,_ N_em_, and N_ti_ = N fertilizer applied at planting, emergence, and tuber initiation, respectively].

**Table 4.** Comparing the performance of the hybrid-LSTM (h-LSTM) and calibrated DSSAT model using the statistical metrics-*nRMSE* and *PR* for individual farms and years while hybrid-LSTM training and testing for estimating SMN daily time series

|  | Farms-years | nRMSE (%) |  | PE (%) |  |
| --- | --- | --- | --- | --- | --- |
|  |  | h-LSTM* | DSSAT | h-LSTM | DSSAT |
| Training | F1-2011 | **13.0** | 17.7 | **8.8** | 12.0 |
|  | F2-2011 | **10.1** | 12.2 | **3.1** | 4.8 |
|  | F3-2011 | **12.8** | 21.4 | **6.2** | 12.5 |
|  | F2-2012 | **9.8** | 19.8 | **1.4** | 6.7 |
|  | F3-2012 | **15.0** | 23.7 | **5.4** | 12.2 |
|  | F4-2012 | 10.0 | **6.1** | 6.6 | **3.8** |
|  | F4-2013 | **11.9** | 16.5 | **7.2** | 11.7 |
|  | F4-2014 | 7.8 | **4.7** | 5.7 | **2.4** |
|  | **Average** | **11.3** | **15.3** | **5.6** | **8.3** |
|  | **Improvement<sup>#</sup> (%)** | **26.1** |  | **32.5** |  |
| Testing-1** | F4-2011 | **7.6** | 9.4 | **2.7** | 3.7 |
|  | F1-2012 | **12.4** | 17.6 | **4.3** | 8.0 |
|  | F1-2013 | 14.5 | **14.4** | 9.8 | **8.6** |
|  | F1-2014 | **12.1** | 15.6 | **10.5** | 14.6 |
|  | **Average** | **11.6** | **14.2** | **6.8** | **8.7** |
|  | **Improvement (%)** | **18.3** |  | **21.8** |  |
| Testing-2*** | F1-2011 | **9.7** | 19.1 | **4.6** | 11.5 |
|  | F2-2011 | **15.0** | 23.5 | **5.4** | 13.2 |
|  | F3-2011 | **16.5** | 26.5 | **8.2** | 15.2 |
|  | F1-2012 | **26.9** | 37.0 | **17.3** | 26.3 |
|  | F2-2012 | **26.5** | 38.7 | **13.9** | 23.5 |
|  | F3-2012 | **28.4** | 40.1 | **15.2** | 25.9 |
|  | F1-2013 | **20.6** | 25.2 | **15.5** | 19.1 |
|  | F1-2014 | **9.1** | 13.9 | **7.1** | 11.9 |
|  | F4-2014 | **11.4** | 11.9 | **8.5** | 8.4 |
|  | **Average** | **18.2** | **26.2** | **10.6** | **17.2** |
|  | **Improvement (%)** | **30.5** |  | **38.3** |  |

\*h-LSTM means the hybrid-LSTM model.
\*\*Testing-1 corresponds to testing results when the hybrid-LSTM was fed with unseen calibrated DSSAT simulated SMN.
\*\*\*Testing-2 corresponds to testing results when the hybrid-LSTM was fed with uncalibrated DSSAT simulated SMN.
<sup>#</sup>Improvement (%) is the percentage improvement shown by the hybrid-LSTM model over the DSSAT model for estimating SMN.

### 3.2. Performance of the hybrid-LSTM model in estimating SMN

Once the calibrated DSSAT model was evaluated, the hybrid-LSTM model was developed using the calibrated DSSAT model simulated SMN and observed SMN samplings for the combination of various fertilizer N rate and timing of application treatments, and farms-years of datasets to improve the SMN estimates. The hybrid-LSTM model performance was then analyzed by feeding both unseen calibrated (section 3.2.1) and uncalibrated DSSAT simulated SMN (section 3.2.2) to determine whether it could improve the calibrated and uncalibrated DSSAT daily SMN simulations using field sampled SMN. Based on the analysis, as the developed hybrid-LSTM model had the knowledge about both the estimates given from the DSSAT model (daily estimates) and the ground truths (sampled estimates), it was able to provide improved estimate of the SMN over DSSAT simulated SMN on daily (Fig. 3, 4) as well as on the sampling days (Fig. 5, Table 4).

**Figure 5.**
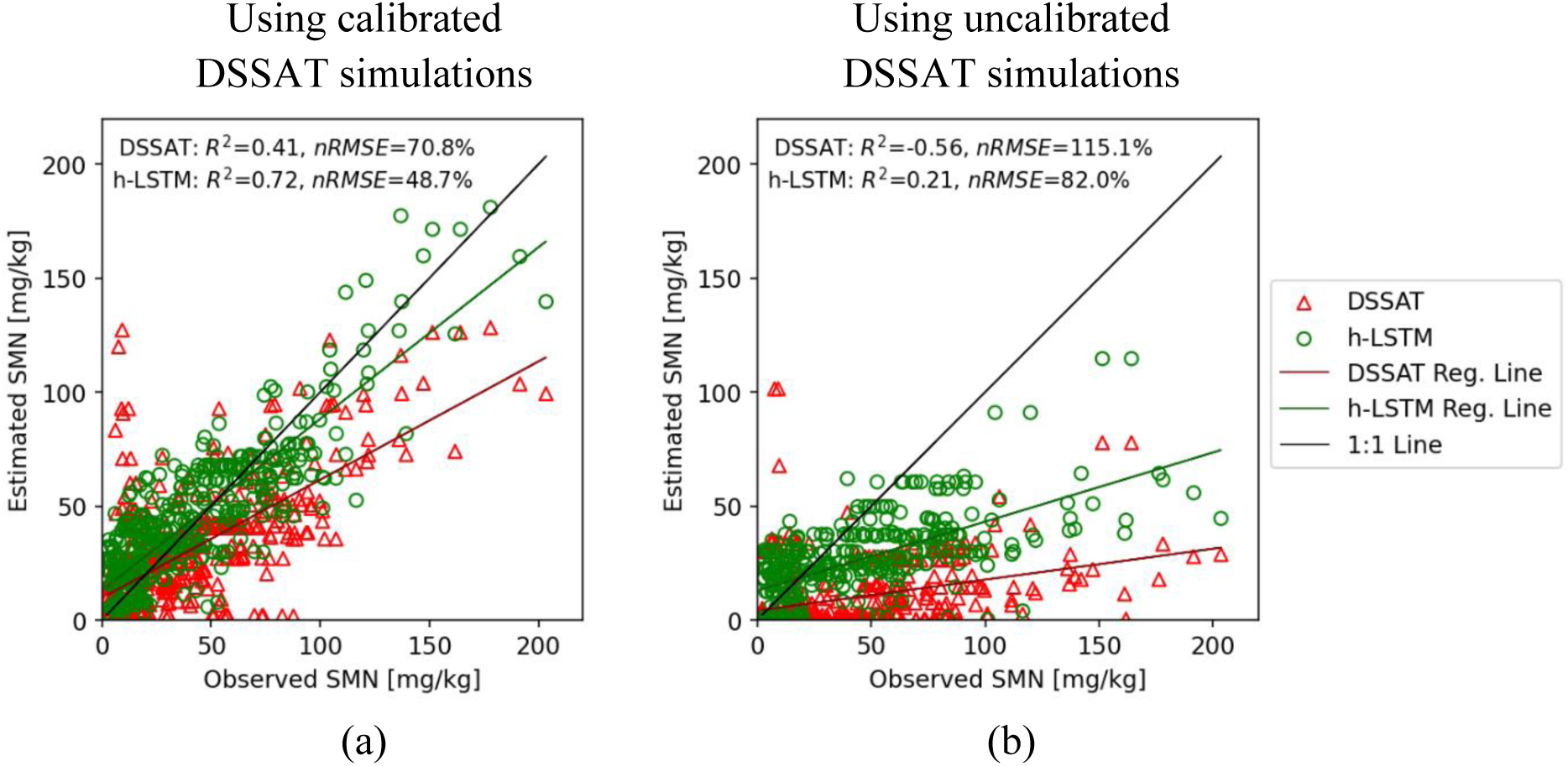
Comparison of the hybrid-LSTM (h-LSTM) estimated and DSSAT simulated SMN against observed SMN at different sampling stages, treatments, and farms-years feeding (a) calibrated and (b) uncalibrated DSSAT simulated SMN in the hybrid-LSTM model. [Reg. = Regression, *R^2^* = Coefficient of Determination, *nRMSE* = Normalized root mean squared error]

#### 3.2.1. The hybrid-LSTM model performance with unseen calibrated DSSAT simulated SMN

Figures 3 and 4 demonstrate the comparison of the SMN values simulated from the hybrid-LSTM and the DSSAT model with observed SMN values for F3-2011 and F1-2013 farm-year combinations, respectively, while training and testing the hybrid-LSTM model (Please see supplementary Fig. S1 and S2 for the results of 2012 and 2014, respectively). Table 4 represents the performance comparison over two metrics of DSSAT simulated SMN and hybrid-LSTM predicted SMN for various farms and years (see section 2.5). As per the results, the hybrid-LSTM model was able to improve the daily SMN estimates over the DSSAT simulated SMN daily values in most of farms-years. The *nRMSE* and *PE* for SMN values estimated using the hybrid-LSTM model [*nRMSE/PE* = 11.3/5.5% (training), *nRMSE/PE* = 11.6/6.8% (testing)] were higher compared to the DSSAT simulated SMN values [*nRMSE/PE* = 12.3/11.3% (training), *nRMSE/PE* = 14.2/11.6% (testing)], both in training and testing the hybrid-LSTM model. Based on these metrics, the hybrid-LSTM model estimated the SMN with 26.1/32.5% (*nRMSE/PE*) and 18.3/21.8% improvement in training and testing farms-years, respectively. In some years farms-years, the *nRMSE* and *PE* of the hybrid-LSTM model increase compared to the DSSAT model. For instance, the *nRMSE/PE* increased from 6.1/3.8% (DSSAT) to 10.0/6.6% (hybrid-LSTM) and 4.7/2.4% (DSSAT) to 7.8/5.7% (hybrid-LSTM), for F4 in 2012 and 2014 used for training the ML model, respectively. Similarly, the *nRMSE* remained almost the same for DSSAT (14.4%) and hybrid-LSTM models (14.5%) while the *PR* increased slightly from 8.6% (DSSAT) to 9.8% (hybrid-LSTM) for F1 in 2013 used for testing the ML model. In all these farms-years (F4-2012, F1-2013, and F4-2014), the hybrid-LSTM model was unable to improve over the DSSAT simulated SMN daily estimates at the sampling stages. Later, the hybrid-LSTM estimated and DSSAT simulated SMN values were compared against those observed on only sampling day for all the farms-years (Fig. 5a). The comparison in Fig. 5a clearly illustrates that the hybrid-LSTM could outperform the DSSAT model in estimating SMN.

### 3.2.2. The hybrid-LSTM model performance with uncalibrated DSSAT simulated SMN

The hybrid-LSTM model was also evaluated for if the model could enhance the uncalibrated DSSAT simulated SMN for different farms-years. These farms-years were similar to what the model was already informed with while training/testing, the only difference being instead of inputting the calibrated DSSAT SMN simulations, the uncalibrated DSSAT SMN simulations were fed into the model. The hybrid-LSTM was able to improve the SMN estimates by 38.3% (*PE*) and 30.5% (*nRMSE*) over uncalibrated DSSAT simulated SMN, as seen in Table 4. The hybrid-LSTM outperformed the DSSAT model in estimating SMN in most farms-years except F4-2014 (please refer to the supplementary Fig. S3 comparing daily SMN estimated using the DSSAT and the hybrid-LSTM models against observed SMN). Figure 5 illustrates a comparison between the hybrid-LSTM estimated and DSSAT simulated SMN against the observed SMN when feeding calibrated (Fig. 5a) and uncalibrated (Fig. 5b) DSSAT simulated SMN in the hybrid-LSTM model for all sampling stages, treatments, and farms-years. With these results, it can be interpreted that the hybrid-LSTM model could improve the DSSAT simulated SMN time-series even when uncalibrated DSSAT model simulated SMN values were fed in the hybrid-LSTM.

### 3.3. Discussion on the overall performance of the hybrid-LSTM model in estimating SMN

Overall, the hybrid-LSTM model outperformed across most farms-years compared to DSSAT. The hybrid-LSTM model improved on the DSSAT simulated SMN daily values regardless of whether the calibrated or uncalibrated DSSAT simulated SMN were fed into it. However, there were a very few exceptions where DSSAT simulations appeared slightly better.

These exceptions could be attributed to the limited SMN observations available for fine-tuning hybrid-LSTM model hyperparameters. Another reason for these exceptions could be due to generalized (same soil properties for all the farms) DSSAT calibration for all the farms which could have led to a distribution shift between both the datasets (DSSAT simulated and the observed SMN data). The generalized DSSAT model did not produce a similar trend for every farm compared to the observed SMN, especially in the case of F1-2011, F3-2012, and F4-2012. In these farms, the DSSAT simulated daily SMN did not align with the patterns observed SMN, despite the DSSAT model sometimes accurately predicted the daily SMN on a few sampling dates. It could be challenging to get a consistent performance with a generalized model in each farm with different soil properties (such as soil textural properties, soil water retention curve, soil organic matter content) (da Silva et al., 2018) under fluctuating water table level and irrigation water distribution in the root zone across the field. In this study, sandy soils were the predominant soil textural class with low soil organic matter. Even a small difference in their texture and organic matter content can change the soil water retention curve affecting the N movement in the soil. da Silva et al. (2018) reported different soil water retention curves due to sand content ranging from 81.9-91.6% in nearby areas. In addition, a great variation in soil moisture is also expected due to the irrigation and drainage cycles during the crop season (Rens et al., 2022). These generalizations in using the DSSAT model generated a trend gap in the datasets (DSSAT simulated vs observed SMN) which might have hampered the overall hybrid-LSTM training. The hybrid-LSTM model trained on such an aberrated dataset would not always give consistent results. Consequently, refining these predictions with fine-tuning might bring the predictions closer to the experimental SMN values for that specific farm-year, but it could affect the overall learning for other farms-years.

Similarly, with cover crop inclusion every year, the soil organic matter from the cover crop residue could also vary among the farms-years and could cause differences in the SMN values with the changes in the net mineralization. Since there was no feature used to track these soil properties variations and cover crop growth dynamics among the farms-years, the hybrid-LSTM model could not incorporate the effect of these changes. Hence, these could be a few of the reasons that the hybrid-LSTM model could not improve the SMN daily estimates over DSSAT simulated SMN while being trained on the generalized DSSAT model simulations. The hybrid-LSTM model outperforms the DSSAT model for F1-2011 and F3-2012, while falling slightly short in the case of F4-2012.

In summary, the hybrid modeling approach incorporating a time-series ML model and a crop simulation model DSSAT proved to be promising in estimating the SMN, especially between the sampling dates. There were a few exceptions and inconsistencies in some farms and years generated due to several reasons mentioned above, could be rectified with more data and improved approaches. In the future, the hybrid model developed could be improved by continuously training on newly available experiment observations from various farms and years. Moreover, encompassing a broader array of scenarios and facilitating a more comprehensive understanding of data patterns and features relationships could further strengthen the modeling approach presented in this article.

## 4. Conclusions

The present study was planned to develop a hybrid ML model to estimate soil mineral nitrogen (SMN) by combining a process-based cropping system model-DSSAT and deep learning-based long short-term memory (LSTM) model in the sub-irrigated potato agroecosystem. The main purpose of the study was to present the novel approach of using the LSTM model with the DSSAT model to estimate the SMN time series in the data-sparse agricultural production system. The hybrid modeling approach could leverage the strengths of the DSSAT model of simulating the daily trend of SMN values and could overcome its weaknesses such as incompetence in replicating a high-water table conditioned sub-irrigated cropping system. The hybrid model developed was able to estimate the SMN time series with a higher accuracy (reducing *nRMSE* from 18 to 30% and *PE* from 21 to 38%) than the DSSAT simulated SMN time series for most of the farms and years. However, there were a few farms and years where the hybrid model could not outperform the DSSAT model. The possible reasons could be the limited experimental observations, incorporating a generalized modeling approach for the farms with slightly different soils, and inconsistencies in the DSSAT simulated SMN on which the hybrid model was initially trained. Overall, the hybrid modeling approach produced a more reliable SMN time series which could aid in developing sustainable N management and provide optimized N fertilizer recommendations minimizing environmental damages for various cropping systems around the world. The proposed modeling strategy could be adapted to estimate the daily time series of other soil nutrients, for instance, phosphorus and sulfur.

## Author Contributions

Rishabh Gupta, Satya K. Pothapragada, Weihuang Xu, Joel B. Harley, Alina Zare, Lincoln Zotarelli: Conceptualization, methodology; Rishabh Gupta, Satya K. Pothapragada: Formal analysis, Investigation, Software; Prateek Kumar Goel, Miguel A. Barrera, Mira Saldanha: Data curation; Rishabh Gupta, Satya K. Pothapragada: Writing—original draft preparation; Joel B. Harley, Alina Zare, Lincoln Zotarelli: Writing—review and editing; Rishabh Gupta: Visualization; Kelly Morgan, Joel B. Harley, Alina Zare, Lincoln Zotarelli: Project administration, funding acquisition. All authors have read and agreed to the published version of the manuscript.

## Acknowledgements

This project was funded by the Florida Department of Agriculture and Consumer Services (Award AWD10728, AWD12838, and AWD15066). The authors acknowledge the support of Judyson de Matos Oliveira, Fernando Rodrigo Bortolozo, and Ayesha Naikodi.

