## Supplementary Document for "Estimating soil mineral nitrogen from data-sparse field experiments using crop model-guided machine learning approach"

**of**

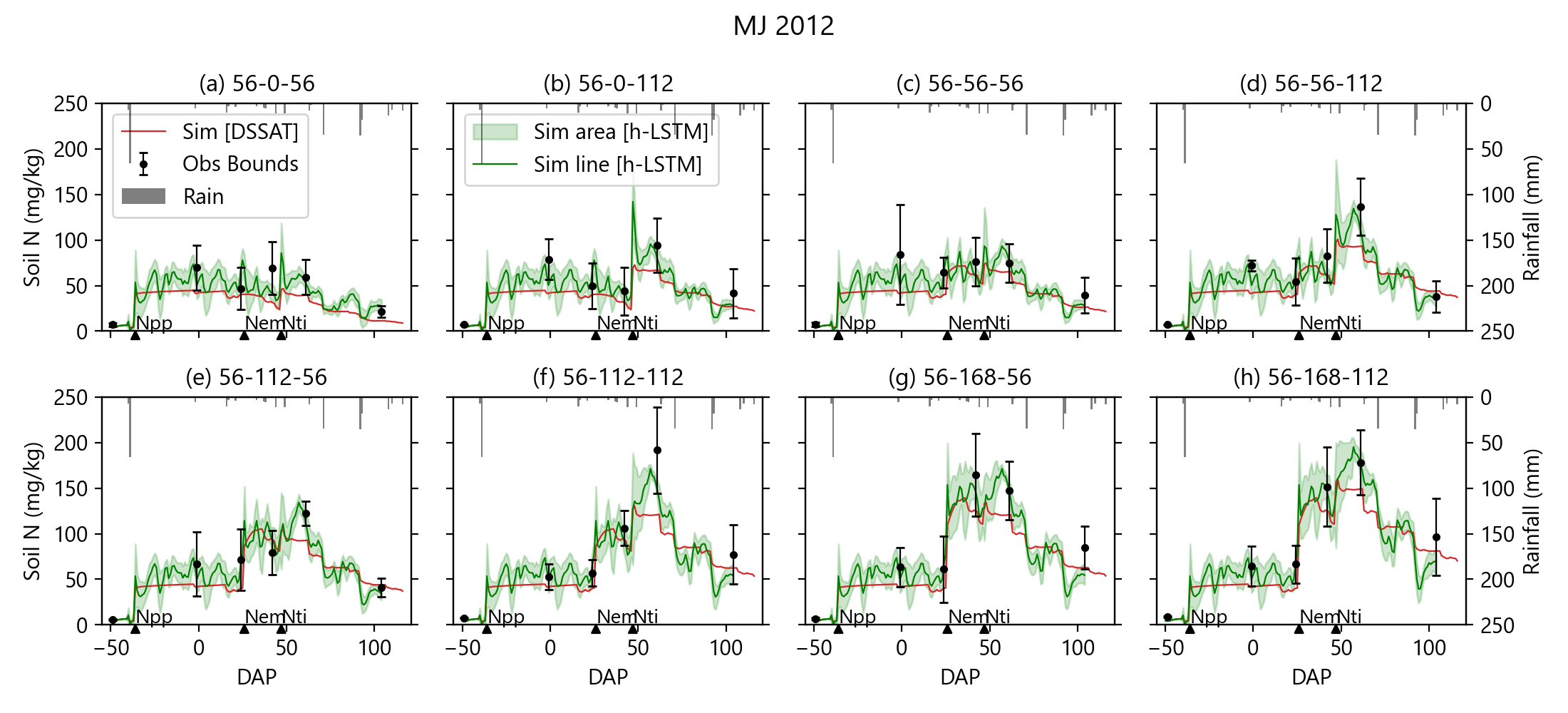

**Figure S1.** Comparison of the observed (black error bars, error bar represents the standard deviation within the replicates), DSSAT simulated (red line) and hybrid-LSTM estimated (green line, green area curve) soil N concentration (0-15 cm) in F2-2012 for different fertilizer N rates and timing of application treatments (subplot title of (a) is 56-0-56 which means N_pp_=56 kg-N ha^-1^, N_em_=0 kg-N ha^-1^, and N_ti_=56 kg-N ha^-1^, the solid triangles below x-axis are the days after planting (DAP) when N fertilizer was applied for N_pp_, N_em_, and N_ti_) while training the hybrid-LSTM model [Sim = Simulated, Obs = Observed; N_pp,_ N_em_, and N_ti_ = N fertilizer applied at planting, emergence, and tuber initiation, respectively].

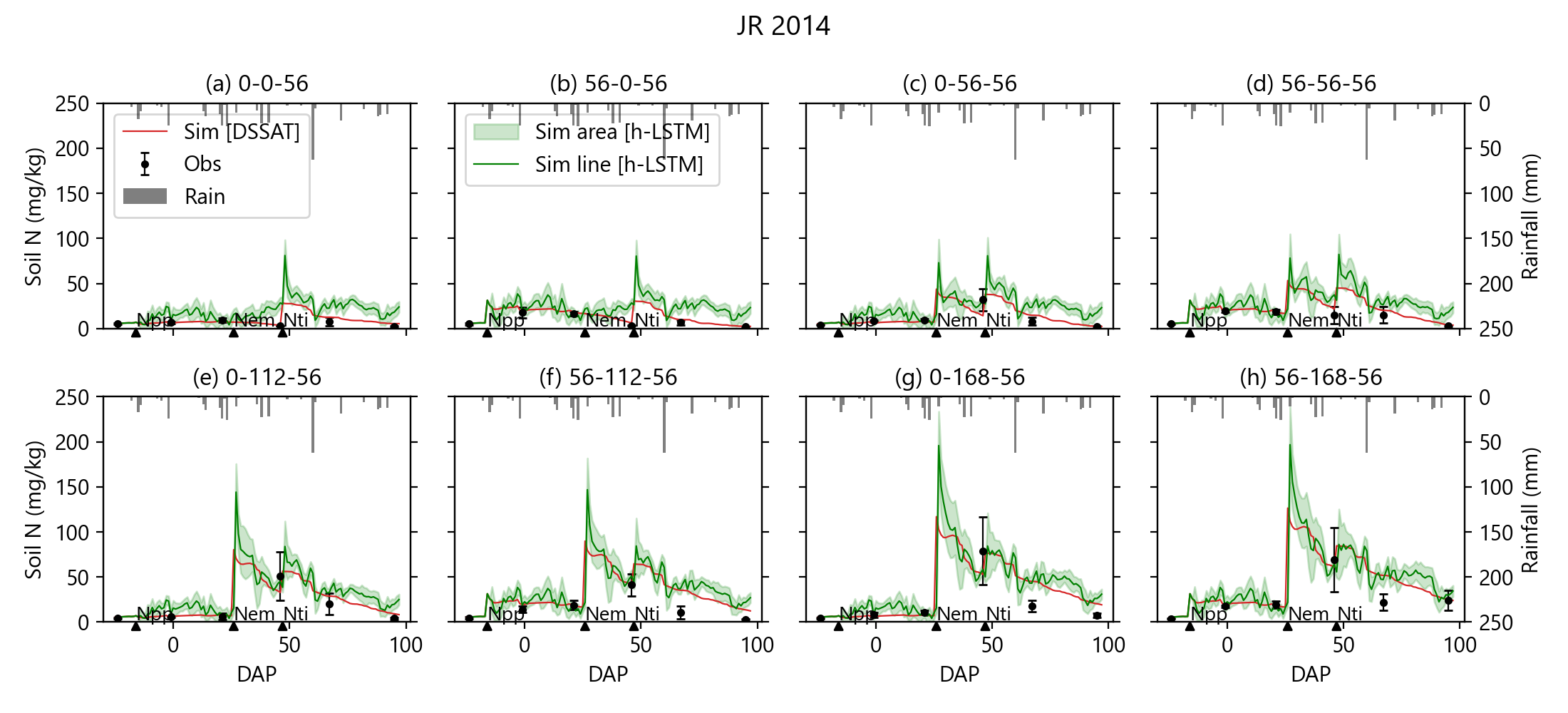

**Figure S2.** Comparison of the observed (black error bars, error bar represents the standard deviation within the replicates), DSSAT simulated (red line) and hybrid-LSTM estimated (green line, green area curve) soil N concentration (0-15 cm) in F4-2014 for different fertilizer N rates and timing of application treatments (subplot title of (a) is 0-0-56 which means N_pp_=0 kg-N ha^-1^, N_em_=0 kg-N ha^-1^, and N_ti_=56 kg-N ha^-1^, the solid triangles below x-axis are the days after planting (DAP) when N fertilizer was applied for N_pp_, N_em_, and N_ti_) while testing the hybrid-LSTM model [Sim = Simulated, Obs = Observed; N_pp,_ N_em_, and N_ti_ = N fertilizer applied at planting, emergence, and tuber initiation, respectively]

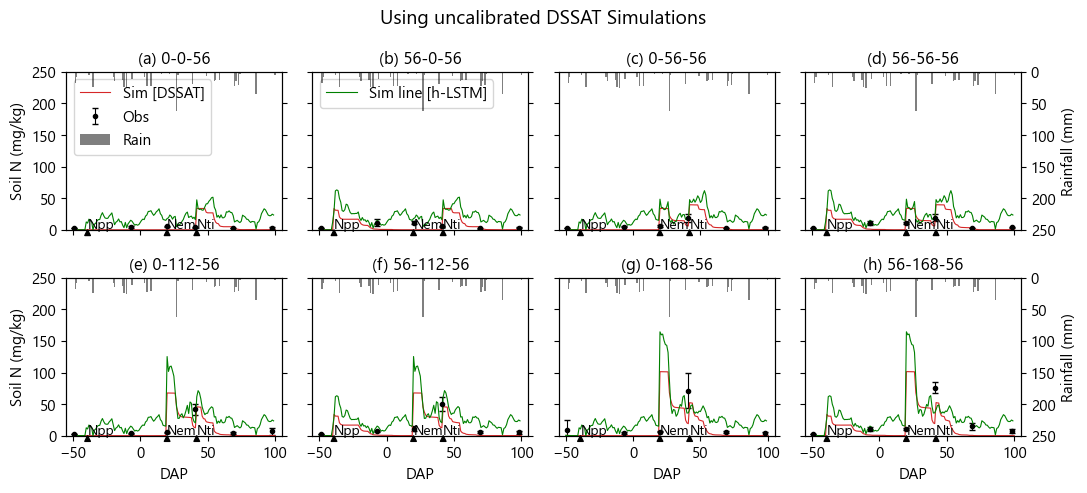

**Figure S3.** Comparison of the observed (black error bars, error bar represents the standard deviation within the replicates), DSSAT simulated (uncalibrated model, shown with red line) and hybrid-LSTM estimated (green line, green area curve) soil N concentration (0-15 cm) in F1-2014 for different fertilizer N rates and timing of application treatments (subplot title of (a) is 56-0-56 which means N_pp_=56 kg-N ha^-1^, N_em_=0 kg-N ha^-1^, and N_ti_=56 kg-N ha^-1^, the solid triangles below x-axis are the days after planting (DAP) when N fertilizer was applied for N_pp_, N_em_, and N_ti_) while testing the hybrid-LSTM model for if it could improve uncalibrated DSSAT simulated SMN [Sim = Simulated, Obs = Observed; N_pp,_ N_em_, and N_ti_ = N fertilizer applied at planting, emergence, and tuber initiation, respectively].
